# Identification and characterization of m6A circular RNA epitranscriptomes

**DOI:** 10.1101/115899

**Authors:** Chan Zhou, Benoit Molinie, Kaveh Daneshvar, Joshua V. Pondick, Jinkai Wang, Nicholas O. Van Wittenberghe, Yi Xing, Cosmas C. Giallourakis, Alan C. Mullen

## Abstract

This study brings together the expanding fields of RNA modifications and circular (circ) RNAs. We find that cells express thousands of m^6^A methylated circRNAs, with cell-type specificity observed between human embryonic stem cells and HeLa cells. m^6^A-circRNAs were identified by RNA sequencing of total RNA following ribosome depletion and m^6^A immunoprecipitation. The presence of m^6^A-circRNAs is corroborated by the identification of complexes between circRNAs and YTHDF1 and YTHDF2, proteins that “read” m^6^A sites in mRNAs.

Furthermore, m^6^A modifications on non-linear RNAs depend on METTL3 and METTL14, the known m^6^A methyltransferase “writer” complex components, suggesting that circRNAs are methylated by the same complexes responsible for m^6^A modification of linear RNAs. Despite sharing m^6^A readers and writers, m^6^A-circRNAs are frequently derived from exons not methylated in mRNAs. Nevertheless, m^6^A-mRNAs that are methylated on the same exons as those composing m^6^A-circRNAs exhibit less stability than other m^6^A-mRNA, and this circRNA-mRNA cross-talk is regulated by YTHDF2. Thus, our results expand the m^6^A regulatory code through identification of the first circRNA epitranscriptome.

## Introduction

RNA base chemical modifications are emerging as a critical layer of post-transcriptional gene regulation. A search of the RNA Modification Database reveals more than 90 RNA modifications found in eukaryotic cells, the vast majority of which are described on heavily modified transfer (t) and ribosomal (r) RNAs (Cantara et al., 2011). However, there is evolving recognition that multiple internal modifications also occur on messenger (m) and long noncoding (lnc) RNAs. N6-methyladenosine (m^6^A) was the first identified mammalian internal mRNA modification and remains the most abundant modification known on mRNAs and lncRNAs (Gilbert and Bell, 2016). A renewed interest in RNA modifications catalyzed by technological advances has revealed the widespread nature of m^6^A in eukaryotic cells from yeast to humans as well as its reversibility in mammalian cells (Dominissini et al., 2012; Jia et al., 2011; Meyer et al., 2012; Schwartz et al., 2013). Furthermore, the identification of proteins that act as “writers,” “readers,” and “erasers” of m^6^A as well as recognition of other internal modifications such as 5- methylcytosine (m5C) (Dubin and Taylor, 1975; Squires et al., 2012), N1-methyladenosine (m1A) (Dominissini et al., 2016; Li et al., 2016; Ozanick et al., 2005), and pseudouridine (Ψ) (Carlile et al., 2014; Lovejoy et al., 2014; Schwartz et al., 2014; Staehelin, 1971) has led to a new field coined ‘epitranscriptomics’. m^6^A has been implicated in all aspects of posttranscriptional RNA metabolism including half-life, splicing, translational efficiency, nuclear export and RNA structure (Lichinchi et al., 2006; Spitale et al., 2015; Wang et al., 2014a, 2015a). At the organismal level, perturbations of m^6^A machinery reveal the functional importance of m^6^A across species including the loss of starvation provoked sporulation in *S. cerevisiae* (Clancy et al., 2002), arrest of embryonic development in *A. thaliana* (Zhong et al., 2008) and *D. melanogaster* (Hongay and Orr-Weaver, 2011), alterations in circadian rhythms (Fustin et al., 2013), as well as a role in the differentiation of mouse and human pluripotent stem cells (Batista et al., 2014; Geula et al., 2015).

The development of m^6^A location analyses utilizing anti-m^6^A antibodies coupled to RNA-sequencing after RNA fragmentation plus (m^6^A-CLIP/ m^6^A-PAR-CLIP) or minus cross-linking (m^6^A-seq/MeRIP-seq) has revealed sites of m^6^A modifications located on thousands of mRNAs and hundreds of lncRNAs (~1 to 3 m^6^A sites per transcript) in numerous primary and transformed cells (Chen et al., 2015; Ke et al., 2015; Linder et al., 2015). Along with site and cell/tissue specificity, m^6^A modifications in resting cells exhibit global enrichment in the 3’UTR near mRNA stop-codons and long internal exons, leading to unique m^6^A-derived transcriptome topology (Dominissini et al., 2012; Meyer et al., 2012). The dynamic and regulated nature of the m^6^A epitranscriptome is highlighted by the observation that certain cellular stresses lead to induction of m^6^A sites in the 5’UTR (Meyer et al., 2015; Zhou et al., 2015). The writing of m^6^A is accomplished via an m^6^A methyltransferase complex composed of a core METTL3 and METTL14 heterodimer (Liu et al., 2014; Wang et al., 2016a, 2016b). Proteins containing the YTH domain directly bind m^6^A sites and act as readers of the m^6^A signal (Wang et al., 2014a, 2015a; Xiao et al., 2016;). YTHDF2 proteins recruit m^6^A-modified mRNAs to nuclear p-bodies promoting RNA degradation (Wang et al., 2014a), while YTHDF1 promotes the translation of m^6^A-modified mRNAs through interaction with translation initiation machinery (Wang et al., 2015a).

The development of m^6^A location analyses utilizing anti-m^6^A antibodies coupled to RNA-sequencing after RNA fragmentation plus (m^6^A-CLIP/ m^6^A-PAR-CLIP) or minus cross-linking (m^6^A-seq/MeRIP-seq) has revealed sites of m^6^A modifications located on thousands of mRNAs and hundreds of lncRNAs (~1 to 3 m^6^A sites per transcript) in numerous primary and transformed cells (Chen et al., 2015; Ke et al., 2015; Linder et al., 2015). Along with site and cell/tissue specificity, m^6^A modifications in resting cells exhibit global enrichment in the 3’UTR near mRNA stop-codons and long internal exons, leading to unique m^6^A-derived transcriptome topology (Dominissini et al., 2012; Meyer et al., 2012). The dynamic and regulated nature of the m^6^A epitranscriptome is highlighted by the observation that certain cellular stresses lead to induction of m^6^A sites in the 5’UTR (Meyer et al., 2015; Zhou et al., 2015). The writing of m^6^A is accomplished via an m^6^A methyltransferase complex composed of a core METTL3 and METTL14 heterodimer (Liu et al., 2014; Wang et al., 2016a, 2016b). Proteins containing the YTH domain directly bind m^6^A sites and act as readers of the m^6^A signal (Wang et al., 2014a, 2015a; Xiao et al., 2016;). YTHDF2 proteins recruit m^6^A-modified mRNAs to nuclear p-bodies promoting RNA degradation (Wang et al., 2014a), while YTHDF1 promotes the translation of m^6^A-modified mRNAs through interaction with translation initiation machinery (Wang et al., 2015a).

We asked whether the concept of an epitranscriptome extends from linear RNAs to circRNAs, defined by the covalent linkage of the 3’ and 5’ ends of spliced RNA transcripts that results in a circularized transcript (Salzman et al., 2012). Back splicing events were initially described in mammalian cells as a source of scrambled exons (Nigro et al., 1991) before these splicing events were linked to circRNAs with the identification of the back splice of *Sry* gene in mice (Capel et al., 1993). Nearly two decades later, the application of high-throughput sequencing of total RNAs depleted of rRNAs revealed that circRNAs are abundant noncoding RNAs (Salzman et al., 2012). Subsequent studies suggested that circRNAs can interact with transcriptional machinery, cyclin-dependent kinases and microRNAs (Hansen et al., 2013; Jiang et al., 2013; Memczak et al., 2013) and can be used as potential biomarkers of various diseases (Cui et al., 2016; Li et al., 2015; Lukiw, 2013; Qin et al., 2015; Shang et al., 2016; Wang et al., 2015b; Xie et al., 2016; Xuan et al., 2016; Zhong et al., 2016). Some circRNAs may also be translated into protein (Wang and Wang, 2015). However, it is unknown if circRNAs are marked by the same m^6^A modification found in mRNAs and lncRNAs.

To test if m^6^A-modified circRNAs exist, we performed m^6^A RNA immunoprecipitation (RIP) on non-fragmented RNA depleted rRNA. Using this approach, we identified the presence of more than one thousand m^6^A-circRNAs in human embryonic stem cells (hESCs). We found that m^6^A-circRNAs were not limited to hESCs and were also present in HeLa cells. Our comparison of m^6^A-circRNA maps between hESCs and HeLa cells revealed both common and cell-type specific m^6^A-circRNA expression patterns. Surprisingly, a large percentage of m^6^A-circRNAs did not overlap with regions containing m^6^A modifications on mRNAs. The m^6^A readers YTHDF1 and YTHDF2 also interact with circRNAs, suggesting that m^6^A modifications on circRNAs are likely to interact with pathways of m^6^A regulation in linear RNAs. In this regard, our analyses uncovered an unexpected connection between m^6^A-circRNAs and mRNA half-life regulated by YTHDF2. Specifically, mRNAs methylated on the same exons that compose m^6^A-circRNAs tend to be less stable than other m^6^A-mRNAs. These results expand our understanding of the breadth of m^6^A modifications and define new regulatory aspects of an m^6^A code through identification of the first circRNA epitranscriptome.

## Results

### RNase R Resistant RNA Species are m^6^A Modified

M^6^A modifications have been described on mRNAs and linear lncRNAs, and we wanted to determine if circRNAs may also be modified by m^6^A. We isolated total RNA from hESCs and performed depletion of rRNA followed by RNase R digestion to degrade linear transcripts **(Figure 1A, top and Figure S1A)** (Suzuki et al., 2006). Anti-m^6^A RNA immunoprecipitation (RIP) was performed after RNase R digestion. Three fractions were analyzed for the presence of m^6^A modifications utilizing anti-m^6^A dot blot: RNase R-treated rRNA-depleted RNA (“input”), the supernatant following m^6^A RIP (m^6^A negative fraction) and the eluate following m^6^A RIP (m^6^A positive fraction) (**Figure 1A bottom**). Bioanalyzer analysis was performed to evaluate the resultant RNA species at each step in this process (Figure S1A). The input RNA, which contains RNase R-resistant circRNA species, is positive for m^6^A modification, as is the eluate fraction, which contains the material immunoprecipitated with the m^6^A antibody. The supernatant, containing the RNA not precipitated with the m^6^A antibody, shows no evidence of m^6^A (**Figure 1A bottom**). It is unlikely that the positive signal in the dot blot in the m^6^A positive (eluate) IP fraction is due to tRNAs. Although tRNAs, like circRNAs, are resistant to RNase R, they have been shown not to be m^6^A modified in mammalian cells (Mishima et al., 2015). In addition, tRNAs are approximately 75 nucleotides in size (Holley et al., 1965), and Bioanalyzer analysis shows loss of the peak that would contain tRNAs following m^6^A RIP (**Figure S1A**, eluate). These results show that RNase R-resistant (nonlinear) RNAs contain a strong m^6^A signal, and suggest that circRNAs contained in this pool may be modified by m^6^A.

**Figure 1.**
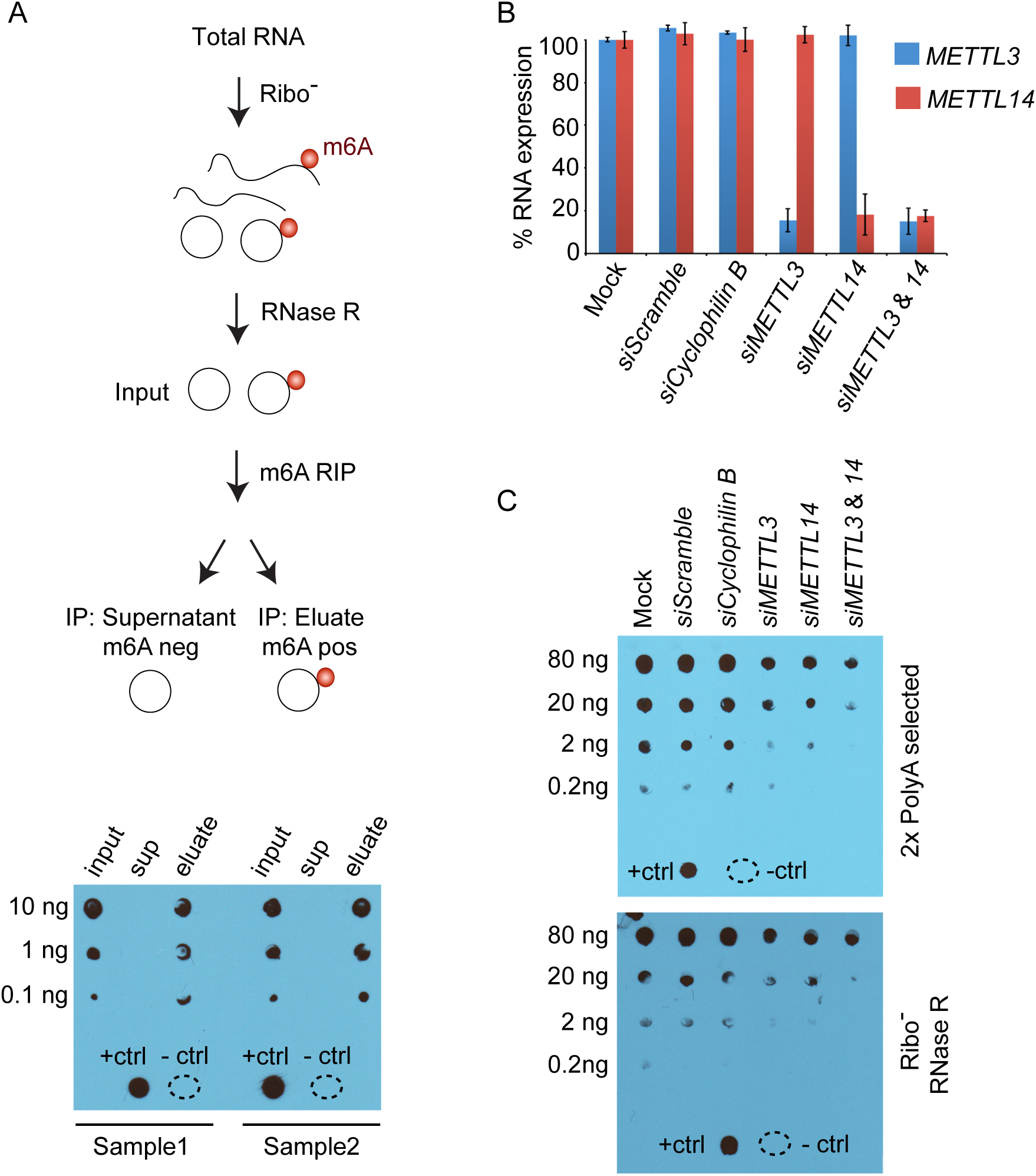
RNase R resistant RNA species are m^6^A-modified. **(A)** The diagram describes how total RNA from hESCs was processed. Dot blots for m^6^A were performed for the indicated amount of RNA. RNA from input (after rRNA-depletion and RNAse R treatment), supernatant (sup) and eluate were probed to detect the m^6^A modification for two replicates (Sample 1 and Sample 2). A positive control (+ctrl) and negative control (-ctrl) containing water are shown at the bottom of the blot. **(B)** 293T cells were transfected with siRNAs that target METTL3 and METTL14 as well as negative controls without siRNA (mock), with scrambled siRNA and with siRNAs that target Cyclophilin B. RNA expression of METTL3 (blue) and METTL14 (red) was normalized to mock transfection. Error bars represent standard deviation. **(C)** Two rounds of polyadenylated RNA selection (**top**) were performed for each siRNA condition in (B). Decreasing amounts of RNA from each condition were probed to detect the m^6^A modification. Controls were performed as in (A). Total RNA was isolated from each siRNA condition in (B). RNA was rRNA-depleted and treated with RNase R to digest linear RNA (Ribo-RNAse R). Decreasing amounts of RNA from each condition were probed to detect the m^6^A modification in circRNAs (**bottom**).

### METTL3 and METTL14 are Required for m^6^A Modifications of Non-linear RNAs

METTL3 and METTL14 physically interact in a synergistic complex, which is required for m^6^A modification of polyadenylated (polyA) RNAs (Liu et al., 2014). The METTL3/METTL14 complex has much higher activity than either isolated protein *in vitro* (Liu et al., 2014), while knockdown of either METTL3 or METTL14 decreases m^6^A levels to a similar degree (Wang et al., 2014b). Therefore, we asked if METTL3 and/or METTL14 are also required for m^6^A modification of non-linear RNAs enriched in circRNAs. We evaluated the impact of depletion of *METTL3* alone, depletion of *METTL14* alone, and the depletion of *METTL3 and METTL14* on m^6^A levels using small interfering (si) RNA-mediated gene silencing. siRNAs targeting of *METTL3* and/or *METTL14* showed efficient knockdown after 48 hrs by qRT-PCR compared to mock, scrambled siRNA and cyclophilin B negative control transfections in HEK-293T cells (**Figure 1B and S1B**). Depletion of *METTL3* and/or *METTL14* resulted in reduced m^6^A methylation of polyA RNAs as expected, based on semi-quantitative m^6^A dot blot analyses (**Figure 1C, top and S1C, top**) (Batista et al., 2014; Liu et al., 2014). We next assayed the effect of siRNA depletion of *METTL3* and/or *METTL14* on rRNA-depleted and RNase R-digested RNA. This analysis demonstrated a reduction in m^6^A modification of non-linear (circRNA-enriched) RNA upon depletion of *METTL3* or *METTL14* individually, but also a synergistic reduction upon combined *METTL3/14* depletion. (**Figure 1C, bottom and S1C, bottom**). Together, these data show that there is an RNase R-resistant (non-linear) fraction of RNA that is dependent on METTL3/14 for m^6^A modification and suggest that circRNAs may be one of these RNA species.

### Embryonic Stem Cells Express m^6^A-modified CircRNAs

To test for the existence of m^6^A-modified circRNAs, we extended the approach of detecting nonlinear back splice junctions to m^6^A RNA fractions in cells. In this regard, we prepared libraries for RNA sequencing and then developed a custom computational pipeline (AutoCirc) to identify back splice junctions in the sequencing data (**Figure 2A and S2A**). We applied our AutoCirc computational pipeline to two biological replicates from hESCs, a cell type for which we have previously published genome-wide m^6^A location analyses (m^6^A-seq) on fragmented polyadenylated RNA (Batista et al., 2014). Sequence data from rRNA-depleted m^6^A positive RNAs were mapped to the human genome. All fully aligned reads, as well as reads that mapped to annotated splice junctions, were discarded. The remaining unaligned reads were used to identify back splice junctions where the 3’ end of a downstream RNA product spliced to the 5’ end of an upstream product. Back splice junctions that contained the flanking GT-AG donor and acceptor sites and junctions that correspond to annotated exon boundaries were considered to form circRNAs (**Figure 2A,** see Materials and Methods).

**Figure 2.**
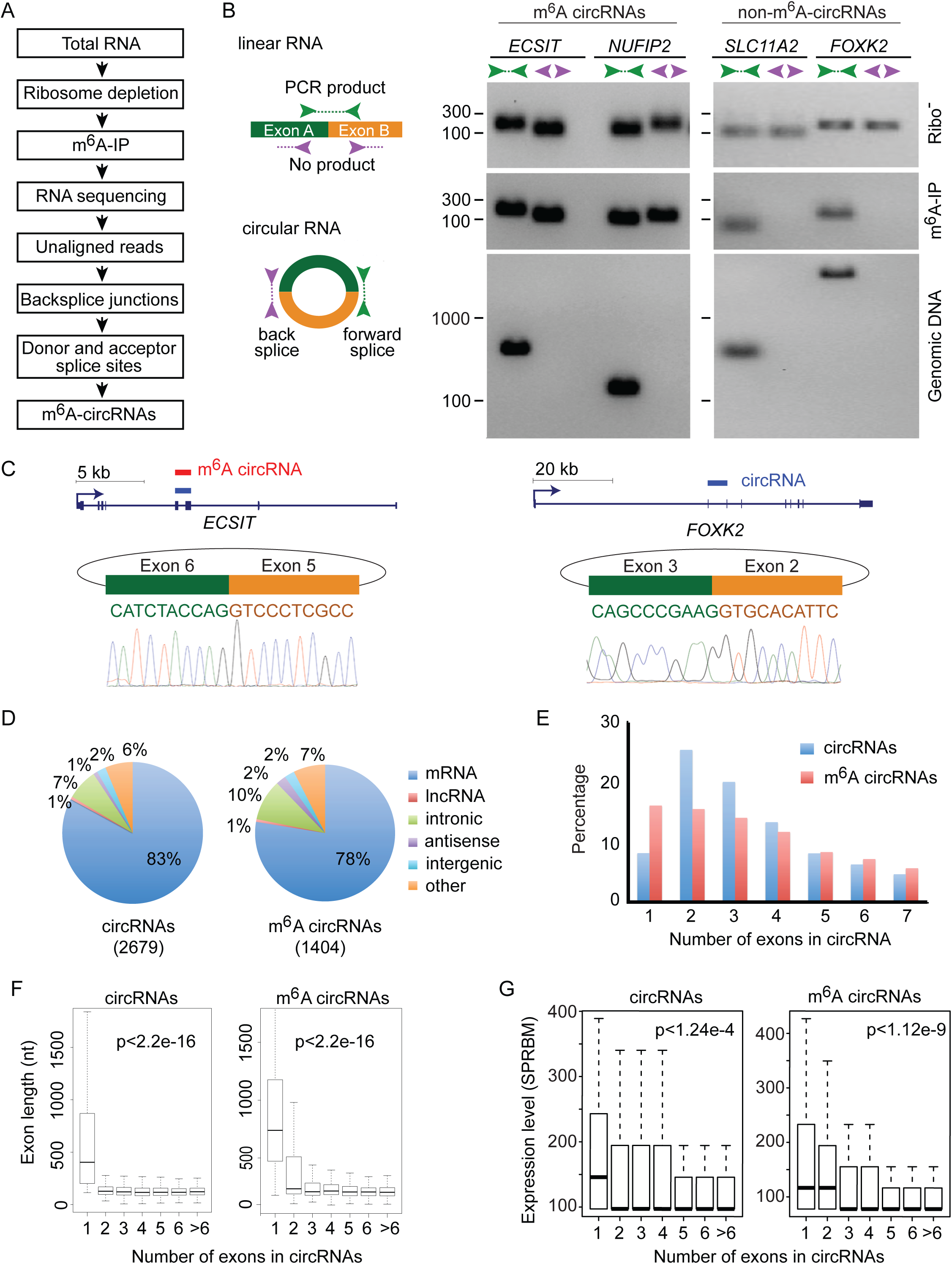
CircRNAs are frequently methylated in human embryonic stem cells. **(A)** Pipeline to identify m^6^A-modifiied circRNAs. **(B)** PCR was performed to detect circRNAs that were methylated (left) and unmethylated (right). Convergent primers (green) were used to amplify forward splice junctions, and divergent primers (purple) were used to amplify back splice junctions produced by circRNAs. PCR was performed on total RNA after rRNA-depletion (Ribo-, top), RNA undergoing rRNA-depletion followed by m^6^A immunoprecipitation (m^6^A-IP, center) and genomic DNA (bottom). Amplicon size is indicated on the left. **(C)** Gene tracks show the locations of total circRNAs (blue box) and m^6^A-circRNAs (red box) within *ECSIT* and *FOXK2.* Rectangles indicate exons and blue lines indicate introns. Arrows represent the start of transcription, and the scale is indicated above each gene. Sanger sequencing results across the back splice junctions are shown below each gene. **(D)** The genomic distribution of circRNAs (left) and m^6^A circRNAs (right) across the genome is indicated. The total number of circRNAs identified in each condition is shown in parenthesis. **(E)** The number of exons spanned by each circRNA (blue) and m^6^A-circRNA (red) was calculated. The percentage of circRNAs (y-axis) spanning up to seven exons (x-axis) is shown. **(F)** The distribution of exon length (y-axis) for circRNAs and m^6^A-circRNAs is plotted based on the number of exons spanned by each circRNA (x-axis). The p values are shown for each condition. **(G)** The expression levels (y-axis) for circRNAs and m^6^A-circRNAs are plotted based on the number of exons spanned by each circRNA (x-axis). Single exon circRNAs are more abundant than circRNAs composed of more than one exon (p<1.24e-4 in total circRNAs and p<1.12e-9 in m^6^A-circRNAs).

The AutoCirc pipeline was then applied to total RNA following rRNA depletion that was not subjected to m^6^A RIP (input) to identify circRNAs that included both m^6^A and non-m^6^A-circRNAs originating from the same sample of hESCs. We identified 2,679 total circRNAs and 1,404 m^6^A-circRNAs by the presence of at least two unique back splice-spanning reads found in the union of biological replicates. We found that only 35% of the circRNAs identified after m^6^A RIP were contained in the pool of total circRNAs (**Figure S2B**), while we would expect the total circRNA pool to contain the m^6^A-circRNAs with sufficient depth of sequencing. We expanded the pool of total circRNAs by using our AutoCirc pipeline to identify circRNAs in non-polyA RNA-seq data of H1 hESCs from the ENCODE project (Consortium, 2012) (see Materials and Methods for details). Over 80% of the m^6^A-circRNAs we identified are contained in the total circRNA pool identified in the two data sets (**Figure S2C**). Reverse transcription (RT)-PCR utilizing convergent and divergent primer sets was then performed to confirm the presence of both m^6^A-circRNAs and non-m^6^A-circRNAs detected by our computational pipeline (**Figure 2B**). Both m^6^A-circRNAs and non-m^6^A-circRNAs were detected by amplification across the back splice junctions in rRNA-depleted samples. However, only the back splice junctions of m^6^A-circRNAs were detected following m^6^A-RIP. These m^6^A- and non-m^6^A-circRNAs were further confirmed by Sanger sequencing across their back splice junctions (**Figure 2C**). These results further verify the accuracy of our computational pipeline and show that many circRNAs contain m^6^A modifications that can be defined by transcriptome-wide sequencing.

We next compared features of total circRNAs and m^6^A-circRNAs. CircRNAs and m^6^A-circRNAs show a similar distribution of genomic localization, where approximately 80% of circRNAs in both categories are derived from exons of protein-coding genes and 1% from exons of IncRNA genes (**Figure 2D**). The majority of circRNAs originate from protein-coding genes span two or three exons, whereas m^6^A-circRNAs are more commonly encoded by single exons (**Figure 2E**). The exons of single exon circRNAs tend to be longer than the exons of multi-exon circRNAs for both total circRNAs and m^6^A-circRNAs (**Figure 2F and S2E**). Single exon circRNAs are also more abundant than multiple-exon circRNAs regardless of methylation status (p<1.24e-4 in total circRNAs and p<1.2e-9 in m^6^A-circRNAs, **Figure 2G**). Furthermore, genes that encode circRNAs and m^6^A-circRNAs are both enriched in the gene ontology (GO) categories of nucleotide binding and ATP-binding activities (**Figure S2F**). Thus, we demonstrate that m^6^A modified circRNAs are widespread (n=1,404 in hESCs) and more likely to be composed of long single exons of genes encoding mRNAs.

During the course of these studies, additional pipelines were developed to identify back splice junctions that define circRNAs. We compared the performance of AutoCirc to CIRCexplorer, which has performed well in other studies (Hansen et al., 2015; Zhang et al., 2014). We found that the two pipelines identify similar numbers of back splice junctions (2,679 for AutoCirc and 2,425 for CIRCexplorer) from rRNA-depleted hESC RNA samples, and 80% of the back splice junctions identified by CIRCexplorer were also identified by AutoCirc (**Figure S2G**). Both AutoCirc and CIRCexlorer also identified a similar low frequency of back splice junction in RNA-seq libraries prepared after polyA selection, which serve as a negative control for circRNAs (**Figure S2H**). These findings suggest that the results from the two pipelines are comparable, whereas our AutoCirc pipeline is (~10 folds) faster and consumes less computing resources (less memory less threads, less number of processes) than CIRCexplorer (**Supplemental Table S1**).

### m^6^A-circRNAs Exhibit Distinct Patterns of m^6^A Modifications Compared to mRNAs

Topologically, m^6^A sites in mRNAs are most common in the last exon (Meyer et al., 2012); however, circularization of the last exon of genes is uncommon (Zhang et al., 2014). Thus, it might be expected that m^6^A-circRNAs would be derived from exons that make up the approximately 20-30% of internal sites of m^6^A modifications on mRNAs. To address this question, we asked if the genes encoding m^6^A-mRNAs are the same genes that encode m^6^A-circRNAs. We found that of 893 genes from which m^6^A-circRNAs are derived, 653 (73%) also encode m^6^A-mRNAs in hESCs (**Figure 3A**). To analyze the overlap in more detail, we examined whether exons methylated in mRNAs are the same exons that are methylated in circRNAs. Surprisingly, the majority (59%) of m^6^A-circRNAs were produced from exons that did not contain m^6^A peaks in mRNAs (**Figure 3B**). Thirty-three percent of m^6^A-circRNAs were produced from genes that encode m^6^A-mRNAs that are methylated on different exons, and 26% of m^6^A-circRNAs were produced from genes that encode mRNAs without detectable m^6^A modification. This observation is also reflected in the different distributions of m^6^A-circRNAs and m^6^A modifications in mRNAs across genes (**Figure 3C**). These results suggest a different set of rules may govern m^6^A-circRNA biogenesis, which will need to be elucidated in the future.

**Figure 3.**
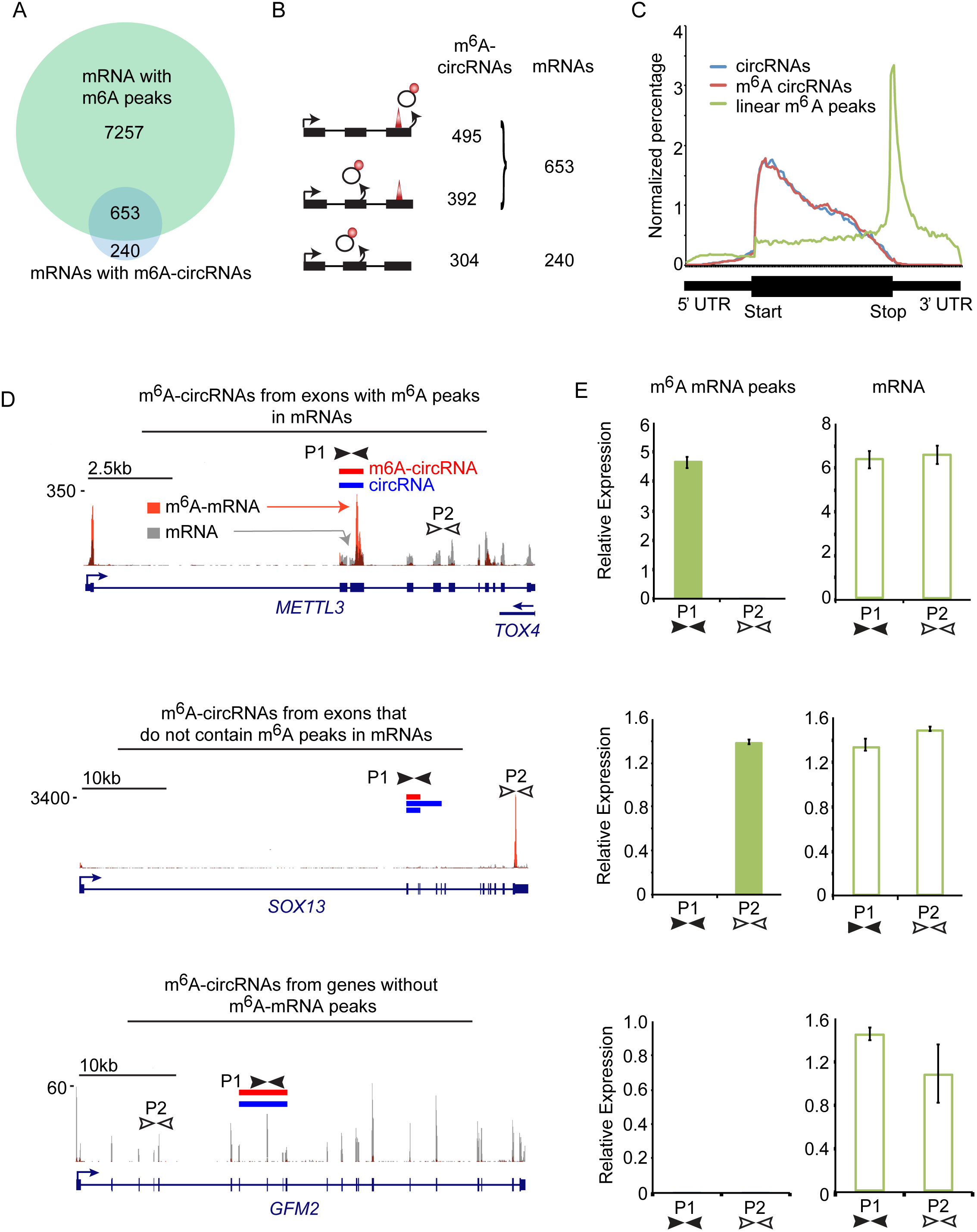
m^6^A-circRNAs are often methylated in regions where m^6^A-mRNAs are not methylated. **(A)** Venn diagram showing the overlap between genes encoding m^6^A-circRNAs and genes encoding m^6^A-mRNAs. The total number of genes in each category is shown. **(B)** The distribution of m^6^A-modified exons between circRNAs and mRNAs: (top) m^6^A sites in the same exons; (middle) m^6^A sites are on different exons from the same parent gene; (bottom) m^6^A sites are only present on circRNAs from a parent gene. The total number of m^6^A-circRNAs and mRNAs are shown. **(C)** Distribution of exons encoding m^6^A-circRNAs (red) and total circRNAs (blue) across compared to the distribution of m^6^A peaks across mRNAs (green). The region from the transcription start site (TSS) to the start of the coding sequence (start) represents the 5’ UTR and the region from the end of the coding sequence (Stop) to the 3’ end represents the 3’ UTR. **(D)** Examples of genes described in (B). m^6^A-circRNAs can be produced from exons that contain m^6^A peaks in mRNAs (top), exons that are different from exons containing m^6^A peaks in mRNAs (middle), or from genes in which no m^6^A peaks are detected in mRNAs (bottom). Red bars indicate m^6^A-circRNAs detected by sequencing. Blue bars represent circRNAs identified by sequencing. Tracks below the circRNAs represent m^6^A peaks from polyadenylated mRNAs before and after m^6^A IP (Batista et al., 2014). Gray peaks represent mRNA background levels, and orange peaks represent sites of m^6^A modification in mRNAs. The locations of primer sets for each transcript are indicated by black and white arrows (P1 and P2). **(E)** qRT-PCR was performed on cDNA prepared following polyA selection and m^6^A IP (m^6^A mRNA peaks) and following polyA selection alone (mRNA) for the genes in (D).

We next performed qRT-PCR on fragmented, polyA-selected RNA before and after m^6^A RIP to confirm the m^6^A status of linear transcripts that also encode circRNAs. The first category represents genes such as *METTL3 and ZNF398* that encode m^6^A-circRNAs from exons that also have an m^6^A peak in mRNAs. Primers were designed to detect the m^6^A mRNA peak within the exons encoding the m^6^A-circRNA (red rectangle, primers P1) and to detect a region of the mRNA without m^6^A enrichment (primers P2) (**Figures 3D, top and S3A, top**). After m^6^A RIP, we detect enrichment of the m^6^A peaks defined by m^6^A-seq (Batista et al., 2014), and there was no amplification from the primers away from the m^6^A peak (**Figures 3E and S3B, top-left)**. Primers set P1 and P2 both amplified fragmented mRNA confirming the presence of both exons in polyA-selected RNA prior to m^6^A IP (**Figures 3E and S3B, top-right)**. These results confirm examples of an m^6^A mRNA peak that is located within exons encoding an m^6^A-circRNA. We next evaluated the second category, representing genes such as *SOX13* and *ARHGEF19,* which encode m^6^A-circRNAs from exons different from those containing m^6^A peaks in mRNA defined by m^6^A-seq. We designed primers to amplify m^6^A-mRNA from exons encoding the m^6^A-circRNA (P1) and primers to amplify the m^6^A-mRNA peak defined by sequencing (P2) **(Figures 3D and S3A, middle)**. We were not able to detect the m^6^A-modification in mRNAs within the exons encoding m^6^A-circRNAs (P1), but we were able to detect m^6^A modification(s) in mRNAs at the site of the m^6^A mRNA peak (P2) (**Figures 3E, and S3B middle**). Both primer sets amplified the mRNA before m^6^A IP, confirming the presence of both exons in the input mRNAs. We then designed primers to confirm the third category of m^6^A-circRNAs encoded by genes such as *GFN2* and *MAPKAP1,* where no m^6^A mRNA peaks were detected in genes encoding m^6^A-circRNAs. Primers were designed to amplify m^6^A-mRNA from exons encoding m^6^A-circRNAs (P1) and another region outside the region encoding m^6^A-circRNAs (P2) (**Figures 3D and S3A, bottom)**. We were not able to detect m^6^A modification(s) in mRNAs within the exons encoding m^6^A-circRNAs or the other regions tested, but both primer sets amplified mRNA, confirming the presence of both exons in the input mRNAs **(Figures 3E and S3B bottom**). These results further validate the finding that numerous m^6^A-circRNAs are produced from exons that do not encode m^6^A peaks in mRNAs.

### m^6^A Methylation of CircRNAs is Cell-type-specific

To determine if m^6^A modifications in hESCs were representative of mammalian cells in general, we sequenced two biological replicates of total RNA from HeLa cells that were depleted of rRNA and then precipitated by m^6^A IP. We identified 854 circRNAs and 987 m^6^A-circRNAs (**Figure S4A**). The genomic distribution, exon length, and number of exons in m^6^A-circRNAs are similar between hESCs and HeLa cells (**Figures S4A-C**). Similar to the distribution of m^6^A-circRNA locations in hESCs, half of the m^6^A-circRNAs identified in HeLa cells originate from exons that do not contain the m^6^A modification in mRNAs (**Figure S4D**).

To determine if m^6^A modifications in hESCs were representative of mammalian cells in general, we sequenced two biological replicates of total RNA from HeLa cells that were depleted of rRNA and then precipitated by m^6^A IP. We identified 854 circRNAs and 987 m^6^A-circRNAs (**Figure S4A**). The genomic distribution, exon length, and number of exons in m^6^A-circRNAs are similar between hESCs and HeLa cells (**Figures S4A-C**). Similar to the distribution of m^6^A-circRNA locations in hESCs, half of the m^6^A-circRNAs identified in HeLa cells originate from exons that do not contain the m^6^A modification in mRNAs (**Figure S4D**).

More than half of the m^6^A-circRNAs detected in HeLa cells were not detected in hESCs (**Figure S4E**), suggesting that many m^6^A-circRNAs are uniquely expressed in the two cell types. HeLa cells and hESCs do not express all of the same genes, so we next asked if the differences of m^6^A-circRNAs between HeLa cells and hESCs could be explained by differences in gene expression. We analyzed circRNAs that were produced only from genes expressed in both HeLa and hESCs and found that 65% of m^6^A-circRNAs detected in HeLa cells were not detected in hESCs even though the parent genes of these circRNAs are expressed in both cell types (**Figure 4A**). When m^6^A-circRNAs are expressed in both cell types, they tend to be expressed at similar levels (**Figure 4B and S4F**). RT-PCR confirmed that unique m^6^A-circRNAs could be detected in the two different cell types. *RASSF8* and *KANK1* are expressed in both HeLa cells and hESCs, yet m^6^A-circRNAs from these two genes are detected only in HeLa cells (**Figure 4C, 4D, top and S4G, S4H, top**). In contrast, *SEC11A* and *TMEFF1* are expressed in both HeLa cells and hESCs, but m^6^A-circRNAs are only detected in hESCs (**Figure 4C, 4D, center and S4G, S4H, center**). m^6^A-circRNAs encoded by *KIF20B* and *NUFIP2* are detected in both HeLa cells and hESCs (**Figure 4C, 4D, bottom and S4G, S4H bottom**). Sanger sequencing confirmed the predicted back splice junctions for m^6^A-circRNAs encoded by *RASSF8, SEC11A,* and *NUFIP2* (**Figure 4E**). These results show that many circRNAs are modified by m^6^A in a cell-type-specific manner.

**Figure 4.**
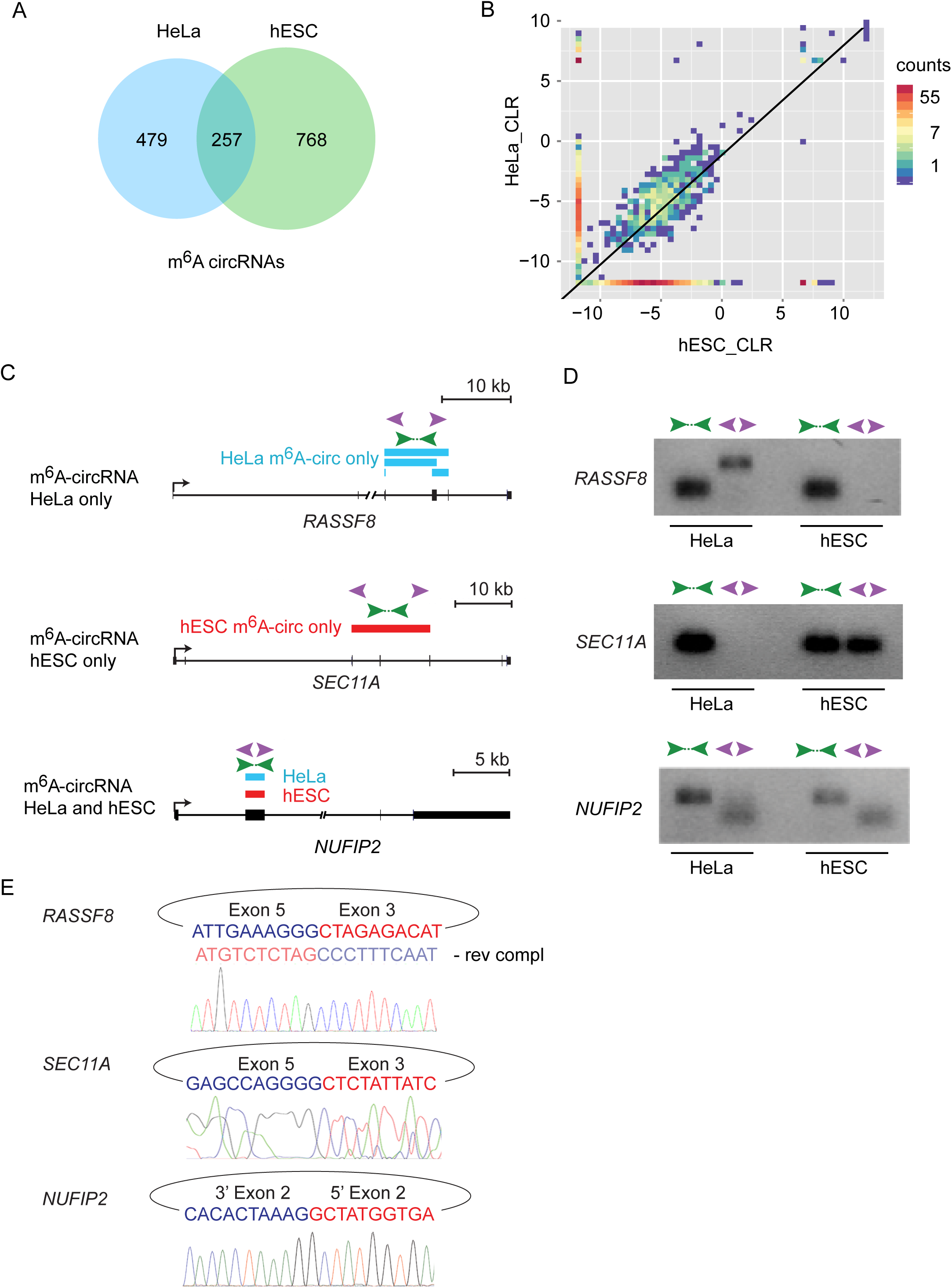
m^6^A-circRNAs show cell-type-specific patterns of expression. **(A)** Venn diagram showing the overlap of m^6^A-circRNAs encoded by genes expressed in HeLa cells and hESCs with RPKM > 1 in both cell types. **(B)** Two-dimensional histogram comparing the expression levels of m^6^A-circRNAs in HeLa cells and hESCs. These m^6^A-circRNAs are encoded by genes, which are expressed in both HeLa cells and hESCs. m^6^A-circRNA expression levels are calculated by the circular-to-linear ratio (CLR) (see Materials and Methods for details). **(C)** Tracks showing examples of m^6^A-circRNAs that that are unique to HeLa cells (top), unique to hESCs (middle), and common between HeLa cells and hESCs (bottom). Blue rectangles indicated m^6^A-circRNAs identified in HeLa cells, and red rectangles indicate m^6^A-circRNAs identified in hESCs. Green arrows indicate the location of primers that amplify across forward splice junctions and purple arrows indicate primers that amplify across back splices. **(D)** RT-PCR was performed on RNA prepared following rRNA depletion, RNase R treatment and m^6^A IP in HeLa and hESC cells, respectively. **(E)** Sanger sequencing was performed on PCR products generated by amplifying across back splice junctions for the indicated genes. Sequencing across the junction is shown in sense for *SEC11A* and *NUFIP2* and antisense (rev compl) for *RASSF8*.

### m^6^A-circRNAs are Recognized by YTHDF1 and YTHDF2 and are Linked to mRNA Stability

YTH-domain family member 1 (YTHDF1) recognizes m^6^A-mRNAs and promotes translation (Wang et al., 2015), while YTHDF2 forms a complex with m^6^A-mRNAs to target RNAs to decay sites (Wang et al., 2014a). We next asked if the YTH domain that recognizes m^6^A-mRNAs also recognizes m^6^A-circRNAs. Ectopic expression of a FLAG-tagged YTHDF1 and YTHDF2 in HeLa cells was used to precipitate the YTHDF1/2 complexes and sequence the bound RNAs (RIP-seq) (Wang et al., 2014a, 2015). We re-analyzed the YTHDF1 and YTHDF2 RIP-seq data using our computational pipeline (**Figure 2A**) to identify circRNAs bound by YTHDF1 and YTHDF2, respectively (**Figure 5A**). We identified 1,155 circRNAs interacting with YTHDF1 and 1,348 circRNAs interacting with YTHDF2 (**Figure 5B**). These circRNAs show a similar distribution across the genome to that of m^6^A-circRNAs (**Figure 5B and Figure S4A**) and are generated from the same categories of genes by GO analysis as m^6^A-circRNAs (**Figure S5B**). In addition, the circRNAs that interact with YTHDF1 and YTHDF2 are formed primarily from exons immediately downstream of the start of coding regions (**Figure 5C**). Twenty-eight percent and 22% of the YTHDF1 and YTHDF2 bound circRNAs, respectively, were also identified as m^6^A-circRNAs in HeLa cells **(Figure 5D)**, and 51% of circRNAs interacting with YTHDF1 also interact with YTHDF2 **(Figure S5A**). To evaluate the possibility that circRNAs may interact with proteins independent of m^6^A modifications, we analyzed RIP-seq data for AGO2 (Polioudakis et al., 2015), a protein which is not known to bind m^6^A modifications. We found that YTHDF1 and YTHDF2-bound circRNAs are significantly enriched in m^6^A-circRNAs compared to AGO2-bound circRNAs (p<0.0041) (**Figure 5E**).

**Figure 5.**
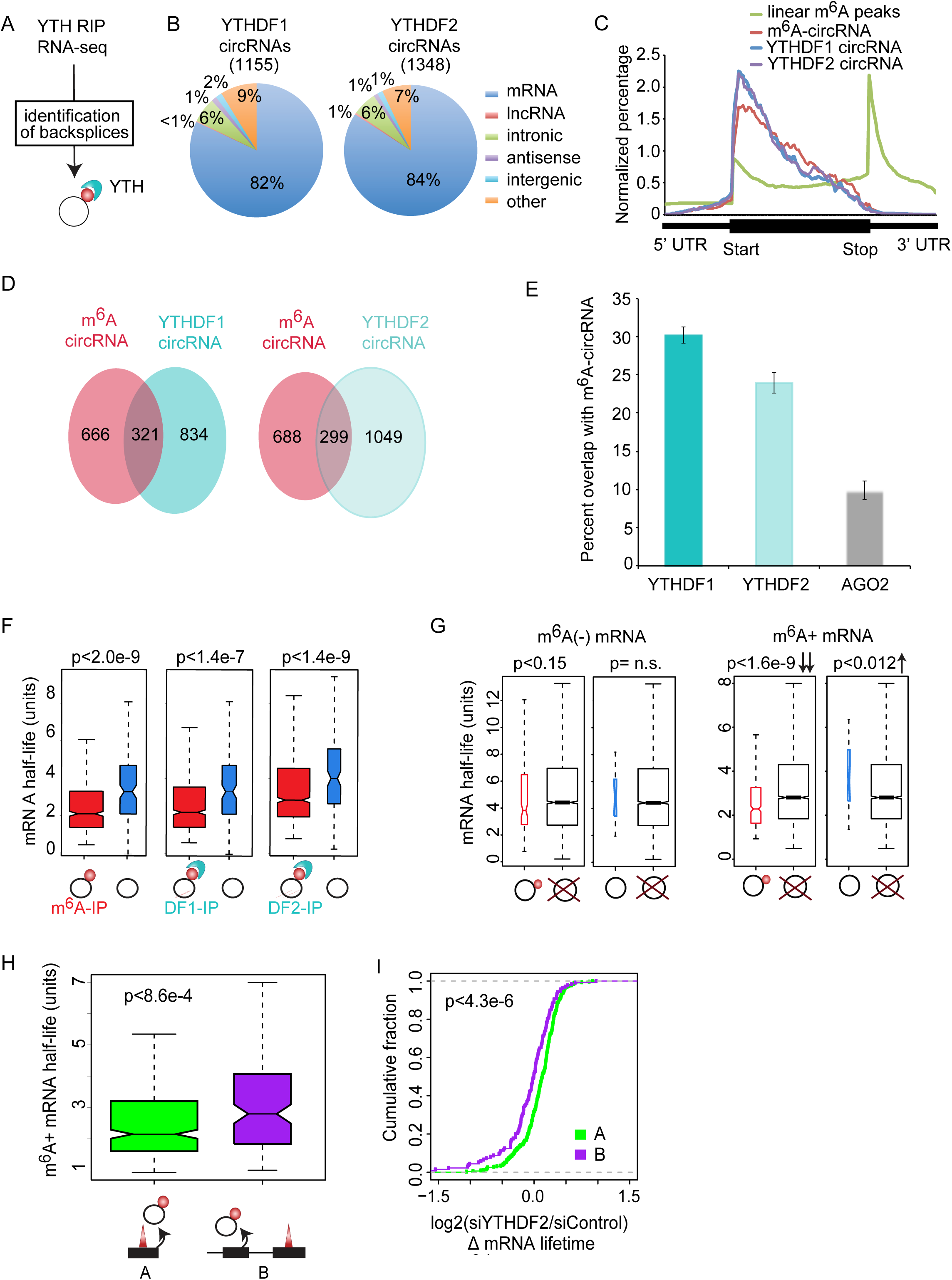
m^6^A-circRNAs bind YTHDF1 and YTHDF2 and identify transcripts with shorter half-lives. (A) Data from YTHDF1 and YTHDF2 RIP-seq (Wang et al., 2014; Wang et., 2015) were used to identify m^6^A-circRNAs in HeLa cells. **(B)** The genomic distribution of circRNAs bound by YTHDF1 and YTHDF2 are shown. The number of circRNAs associated with each protein is indicated in parenthesis. **(C)** The distribution of exons encoding circRNAs associated with YTHDF1 (blue) and YTHDF2 (purple) compared to exons encoding m^6^A-circRNAs (red) and the distribution of m^6^A peaks across mRNAs (green) is shown. **(D)** Venn diagram shows the overlap between m^6^A-circRNAs identified in HeLa cells and m^6^A-circRNAs bound by YTHDF1 (left) and YTHDF2 (right). **(E)** The percentage of m^6^A-circRNAs identified by m^6^A IP in HeLa cells and bound by YTHDF1, YTHDF2, or AGO2 are shown. Error bars represent standard deviation. **(F)** The mRNAs half-life for parent genes that encode m^6^A-circRNAs (left), YTHDF1-bound (DF1-IP) circRNAs (middle), or YTHDF2-bound (DF2-IP) circRNAs (right). Black rings represent circRNAs, red circles indicate m^6^A modification, and light blue structures represent YTH proteins. **(G)** The mRNA half-life for m^6^A negative (-) (left) and m^6^A positive (+) (right) mRNAs whose parent genes encode m^6^A-circRNAs (black ring with red circle attached), circRNAs without m^6^A (black ring), or no circRNAs (black ring with red "X”). The p values are indicated above each comparison and indicate statistically significant changes in half-life. **(H)** The half-life of m^6^A-mRNAs whose parent genes also encode m^6^A-circRNAs from the same exon(s) where an m^6^A peak is found in mRNA (labeled A) is compared to the half-life of m^6^A-mRNA whose parent gene encodes an m^6^A circRNA at exon(s) where no m^6^A peak is found in mRNAs (labeled B). The p value is indicated. **(I)** The change in half-life (log2(siYTHDF2/siControl)) was calculated after depletion of YTHDF2 as described (Wang et al., 2014a) for condition A (green) and condition B (purple). The p values were calculated using Wilcoxon-Mann-Whitney test. (Supplemental Figure 6 shows the results for the second replicate).

YTHDF2 interacts with m^6^A-mRNAs to regulate mRNA stability (Wang et al., 2014a), and we asked if the presence of m^6^A-circRNAs was connected to the half-life of m^6^A-mRNAs. We found that mRNAs encoded by the parent genes of m^6^A-circRNAs also have a shorter half-life than mRNAs encoded by the parent genes of non-m^6^A-circRNAs (**Figures 5F and S5C**). This finding is observed regardless of whether the mRNA contains an m^6^A modification or not. This result is observed for m^6^A-circRNAs identified by m^6^A IP, interaction with YTHDF1, and interaction with YTHDF2. We next performed the same analysis but separated m^6^A negative (labeled as m^6^A(-)) from m^6^A positive (labeled m^6^A(+)) mRNAs (**Figure 5G and S5D**). This analysis shows that m^6^A-mRNAs encoded by the parent genes of m^6^A-circRNAs are the only group with significantly reduced half-lives compared to genes not encoding any circRNAs. These results suggest that in addition to the m^6^A modification of mRNAs being associated with a shorter mRNA half-life (Wang et al., 2014a), those m^6^A-mRNAs encoded by the parent genes of m^6^A-circRNAs have shorter half-lives among all m^6^A-mRNAs.

Genes encoding both m^6^A-mRNAs and m^6^A-circRNAs can produce these RNA products either from the same exons (**Figure 5H, diagram A**) or different exons (**diagram B**). The m^6^A-mRNAs with the shortest half-lives are those in which m^6^A-circRNAs are produced from the same exons that have m^6^A modifications in mRNAs (**Figure 5H**). Because the m^6^A modification is most commonly enriched in the 3’ UTR of mRNAs (Batista et al., 2014; Meyer et al., 2012) while circRNAs are enriched in the gene body (**Figure 3C and 5C**), we next examined if differences in the location of the m^6^A modification in mRNAs could explain the difference in half-lives. We find that there is no significant difference in the half-lives of m^6^A-mRNAs that are modified in the gene body or 3’ UTR (**Figure S5F**). Finally, we find that m^6^A-mRNAs that are methylated in the same exons that are methylated in circRNAs show increased half-lives with depletion of YTHDF2 (**Figures 5I and S5G**), suggesting that stability of this subset of RNAs is controlled by YTHDF2 through binding the m^6^A-circRNAs.

### Discussion

This study brings together two rapidly expanding fields: RNA modifications and circRNAs. Here, we describe the first circRNA epitranscriptome by identifying the presence of thousands of m^6^A-circRNAs. m^6^A-circRNAs are recognized by YTHDF1 and YTHDF2, as are m^6^A-mRNAs and may also be methylated by METTL3 and METTL14, the same methyltransferases responsible for the m^6^A modification in mRNAs. However, we find that many m^6^A-circRNAs originate from regions where m^6^A modifications are absent on mRNAs. These results indicate that the writing and reading machineries of m^6^A modification are similar in both mRNAs and circRNAs, but can produce different patterns of m^6^A modifications in different RNA products. Furthermore, we identify many m^6^A-circRNAs expressed in a cell-type-specific pattern even when their parental genes are expressed in both cell types, suggesting that the m^6^A modification of circRNAs may be regulated to control different biological functions between different cell types.

The discovery of m^6^A-circRNAs also raises many new questions that will need to be addressed in future studies, including the significance of m^6^A-modifications on exons that compose circRNAs but are not modified in mRNAs. Nevertheless, our results show that identification of m^6^A-circRNA patterns may be useful to identify different cell types/states even in the absence of significant changes in baseline mRNA expression, leading to a new methodology to fingerprint cells. Furthermore, it will be of interest to address whether m^6^A-circRNAs are found in extracellular vesicles which have recently been identified to contain circRNAs and proposed as a mechanism to clear circRNAs from cells (Lasda and Parker, 2016).

In terms of m^6^A-circRNA functionality, we provide evidence of cross-talk between m^6^A-modified mRNAs and circRNAs that affects mRNA half-life in a YTHDF2 dependent manner. We can only speculate on the mechanism for now. However, one potential model is that m^6^A-circRNAs and m^6^A-mRNAs encoded by the same exons are bundled as part of a chromatin-associated liquid phase transition leading to a nuclear “liquid droplet” (Caudron-Herger et al., 2016; Marzahn et al., 2016) and continue to be a topologically and organizationally distinct information packets. Whereas m^6^A-circRNAs arising from non-methylated exons of mRNAs are not bundled with mRNAs or are contained in other bundles. It is possible that these information packet(s) can be transmitted to the cytosol leading to differential mRNA processing via interaction with cytosolic liquid droplets, that may include m^6^A binding to YTH domain proteins, which harbor poly Q unstructured domains (Guo and Shorter, 2015; Wang et al., 2014a; Zhang et al., 2015). We postulate that circRNAs in general may exhibit unique tuning qualities on liquid droplets, affecting surface tension, stability, size and/or longevity. m^6^A-circRNAs may further modify these characteristics given their ability to interact with YTH/PolyQ proteins as well as other RNA binding proteins (Guo and Shorter, 2015; Lin et al., 2015).

Controlling the state of m^6^A modifications on circRNAs may act as switches to control circRNA functionality. For example, the presence of m^6^A modifications beckons the question as to whether m^6^A-circRNAs can be translated. This possibility arises given that YTHDF1 recognizes m^6^A-mRNA to promote translation via recruitment of translation initiation factors, m^6^A itself has been shown to recruit initiation factors resulting in cap-independent translation (Meyer et al., 2015; Zhou et al., 2015), and circRNAs can be engineered to be translatable with internal ribosome entry sites (Chen and Sarnow, 1995; Wang and Wang, 2015). Thus, it is possible that the m^6^A modification could convert non-coding circRNAs to coding circRNAs.

We find that many m^6^A-circRNAs originate from regions where m^6^A modifications are absent on mRNAs, suggesting that while the writing and reading machineries of m^6^A modification are similar in both mRNAs and circRNAs, different patterns of m^6^A modifications are produced in different RNA products. How do we reconcile the presence of m^6^A-circRNAs arising from exons that are non-methylated in polyA-selected mRNA species? This may stem from the presence of m^6^A on such exons in pre-mRNA that are retained with circularization, while the site is removed from mRNAs by m^6^A demethylases such as FTO in mRNAs. Alternatively, pre-mRNAs containing m^6^A modifications at certain locations are not demethylated and instead are rapidly degraded if they do not become circRNAs. This process would prevent detection during standard polyA selected m^6^A-seq. A third possibility is that circRNAs are methylated after transcription in the nucleus or cytosol. In this scenario circRNA production and mRNA methylation are separated in space and time. A fourth is the possibility is that circRNA methylation is linked to the process of circRNA formation. This stems from the observation that introns flanking circRNAs in mammalians are enriched in transposable elements (TEs), including Alu repeats (Jeck et al., 2013; Zhang et al., 2014), to facilitate the formation of stem loops during back splicing events.

In summary, we present transcriptome-wide identification of m^6^A-circRNAs, extending the concept of the RNA epitranscriptome to circRNAs. We provide evidence that m^6^A modifications to circRNAs are written and read by the same machinery used for mRNAs, but often at different locations, and we implicate m^6^A-circRNAs in mRNA stability. Our results establish a fertile area of new investigation to define the breadth and function of covalent modifications in circRNAs.

## Experimental Procedures

### RNA isolation, m^6^A immunoprecipitation, and library preparation

Total RNA was obtained by TRIzol extraction followed by DNAse I treatment. Ribosomal RNA (rRNA) depletion was performed twice using the Invitrogen RiboMinus Kit starting with 10 μg of total RNA. For RNAse R treatment, 5 μg of rRNA-depleted total RNA was treated with 5 units of RNAse R per μg twice. 100 ng of RNA was used for library construction for RNA-seq with modifications to the Illumina TrueSeq Stranded mRNA Sample Preparation Guide as detailed in Supplementary Experimental Procedures. PolyA RNA selection was performed twice using the Dynabeads mRNA Purification Kit with 7.5 μg of total RNA input. Either 20 μl 2X RNase R treated RNA or 2x RiboMinus-treated RNA were used as input for m^6^A-IP using an anti-m^6^A antibody obtained from Synaptic Systems (cat.# 202 003). Please see Supplemental Experimental Procedures for additional details.

### m^6^A dot blots

RNA was allowed to bind to an Amersham Hybond membrane by gravity prior to UV crosslinking. The membrane was incubated with a rabbit polyclonal antibody recognizing m^6^A followed by an anti-rabbit secondary antibody before visualization on film using ECL Western Blotting Substrate. Please see Supplemental Experimental Procedures for additional details.

### Computational pipeline for detecting circRNAs

We performed directional, 100 × 100 paired-end sequencing for all our libraries. The paired reads were treated as independent reads and mapped to the human reference genome (hg19). We used Bowtie2 (Langmead and Salzberg, 2012) to identify and discard all sequences that mapped to a contiguous region of genomic DNA. We then developed our computational pipeline (AutoCirc) in C++ to scan the 20 nucleotides at both ends of each 100 nt sequence of unmapped reads to identify back splice junctions. Back splice junctions that contained the canonical GT-AG splice donor and acceptor sequences in the flanking intron boundaries or that occurred at known exon boundaries were considered to represent circular RNAs. CircRNA expression was quantified by spliced reads per billion unique mapped reads (SRPBM) (Jeck et al., 2013). We applied the same pipeline to identify the circRNAs from all our samples. Please see Supplemental Experimental Procedures for additional details. CircRNAs identified in this study are contained in Supplemental Table S2. We obtained two replicates of H1 hESC long polyA minus RNA-seq data from the ENCODE project: http://hgdownload.cse.UCSC.edu/goldenPath/hg19/encodeDCC/wgEncodeCshlLongRnaSeq and the polyA selected RNA-seq data from GEO (acc: GSE41009).

Our AutoCirc pipeline and CIRCexplorer pipeline were applied to identify the circRNAs from all hESC samples and m^6^A**-** circRNAs from hESC eluate samples.

### Comparison of the expression levels of m^6^A circRNAs between hESCs and HeLa cells

To evaluate the relative abundance of circRNAs to their cognate mRNAs, we calculated the circular-to-linear RNA ratio (CLR), computed as (log2(back splice reads/maximum of the reads covering the two end junctions of linear splice sites) (Rybak-Wolf et al., 2015). We then used CLR values to compare the expression levels of m^6^A circRNAs between hESCs and HeLa cells.

### Density distribution of circRNAs and m^6^A-peaks across the entire genes

All parent genes of circRNAs were divided into 50 bins representing the 5’ UTR, 100 bins representing the coding DNA sequence (CDS), and 50 bins representing the 3’ UTR. The density distribution of each circRNA across each bin was calculated and normalized to the size of circRNAs and the total number of circRNAs from each sample. The same approach was adopted to calculate the density distribution of m^6^A-peaks across methylated mRNAs as previously defined (Batista et al., 2014; Wang et al., 2014a).

### Identification of circRNAs interacting with YTHDF1, YTHDF2, and AGO2

We downloaded the two replicates of single-end RNA-seq data for YTHDF1 and YTHDF2 RIP from HeLa cells from GEO (GSE63591 and GSE49339). Four replicates of paired-end RNA-seq data from AGO2 RIP in HeLa cells were obtained from GEO (GSE64615). We applied our computational pipeline to identify circRNAs from these RNA-seq data. We treated four replicates of paired-end RNA-seq data of AGO2 RIP as eight replicates of single-end RNA-seq data. The circRNAs identified by our AutoCirc pipeline with at least two reads in the all replicates are considered as potential circRNAs pulled from RIP. We counted the circRNAs with a single read or greater as present in a replicate as long as there were at least two reads supporting a specific back splice in the union of all replicates. Error-bars in Figure 5E are calculated based on the number of circRNAs present in each replicate.

### Evaluation of mRNA half-life

We obtained mRNA half-life data from siControl and siYTHDF2 in HeLa cells (Wang et al., 2014a). We separated the mRNAs into the following groups: (a) mRNAs produced by parent genes of m^6^A-circRNAs; (b) mRNAs produced by parent genes of non-m^6^A-circRNAs; (c) mRNAs produced by parent genes of circRNAs bound by YTHDF1; (d) mRNAs produced by parent genes circRNAs bound by YTHDF2. We further subdivided these mRNAs groups into mRNAs that contain m^6^A modifications and mRNAs that do not contain m^6^A modifications (Wang et al., 2014a). mRNAs produced by genes that did not produce circRNAs were used as a control group. We also compared the half-lives of m^6^A-mRNAs methylated in the same exons as m^6^A-circRNAs to the half-lives of m^6^A-mRNAs methylated in different exons from m^6^A-circRNAs encoded by the same gene. The analyses above were performed using siControl samples. To examine how the interaction between YTHDF2 and m^6^A-circRNAs affects m^6^A-mRNA half-life, we plotted the accumulation fraction curve of the log2-transformed changed half-life between siYTHDF2 and siControl cells for m^6^A-mRNAs methylated in the same exons as m^6^A-circRNAs and m^6^A-mRNAs methylated in different exons from m^6^A-circRNAs encoded by the same gene.

### Data access

Raw RNA-seq data produced for this study and processed data including coordinates of circRNAs in BED and read density in BigWig format are available in the GEO (GSE85324). *The following link has been created to allow review of record GSE85324 while it remains in private status:*

https://www.ncbi.nlm.nih.gov/geo/query/acc.cgi?token=mtufwuekbdmdhaz&acc=GSE85324

## Supplemental Information

Supplemental information includes Extended Experimental Procedures, five supplemental figures and three supplemental tables.

## Author Contributions

C.C.G. and A.C.M. conceived the study. C.C.G. and A.C.M. designed experiments with C.Z. and B.M. Computational analyses were performed by C.Z. with support from J.V.P., J.W. and Y.X. Bench experiments were performed by B.M. with assistance from K.D. and J.V.P. The manuscript was written by C.Z., C.C.G. and A.C.M. with input from B.M.

## Acknowledgements

We would like to thank P. Dedon and K. Jeffrey for helpful discussions, and thank the MGH Sequencing core and Tufts University Genomics core for RNA sequencing. This work was supported National Institutes of Health (NIH) grant R01GM088342, an Eli & Edythe Broad Center of Regenerative Medicine and Stem Cell Research at UCLA and Rose Hills Foundation Research Award, and an Alfred Sloan Research Fellowship (Y.X.). This work was also supported by Massachusetts General Hospital start-up funds (C.C.G. and A.C.M.).

---

**Figure S1, related to Figure 1:**
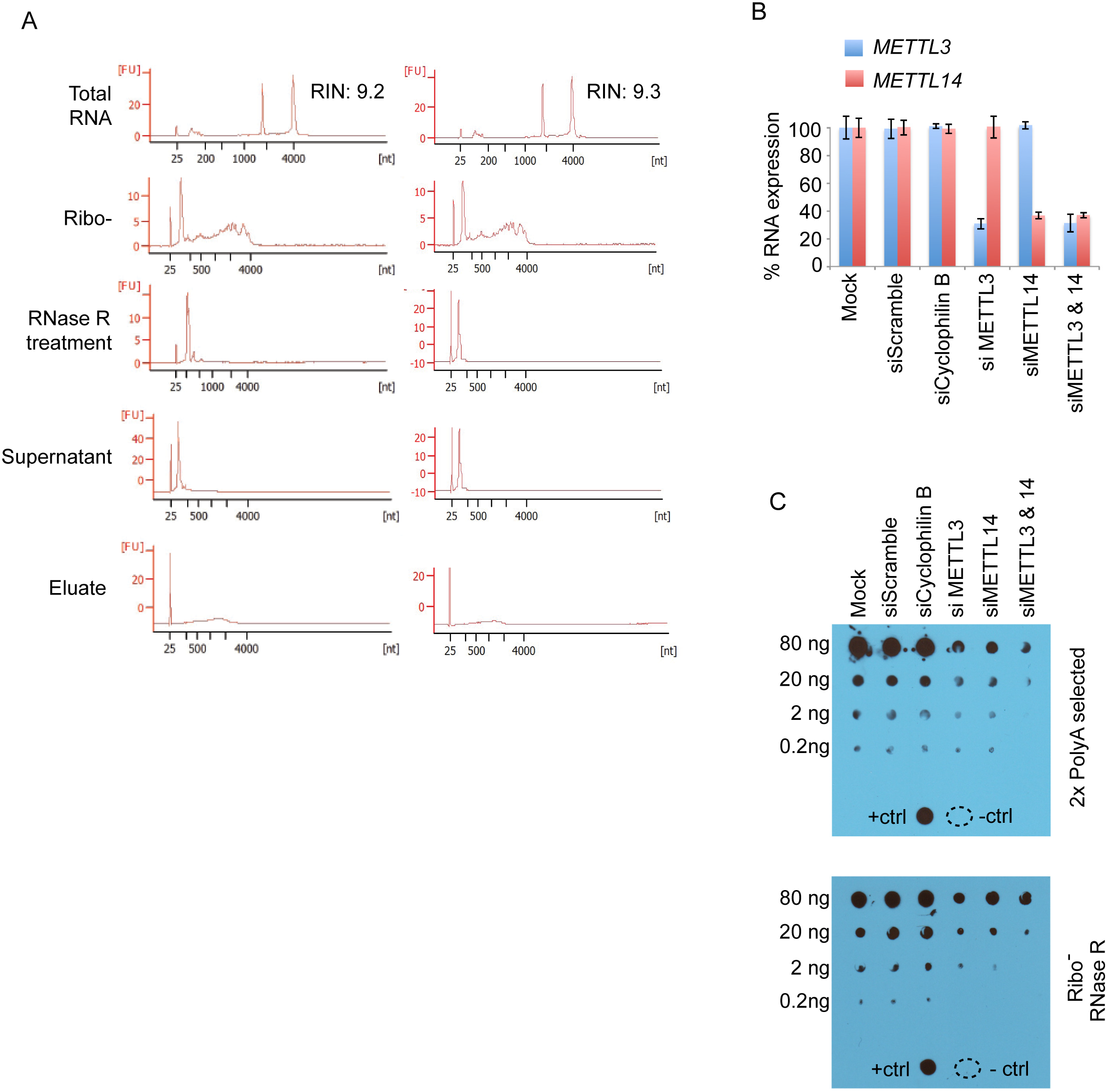
RNase R resistant RNA species are m6A modified. **(A)** Bioanalyzer analysis of RNA samples processed for dot blot (Fig 1A). RIN scores are indicated for total RNA for each replicate. Eluate refers to RNA precipitated by m^6^A IP and supernatant is the RNA pool that was not precipitated by m^6^A IP. **(B)** 293T cells were transfected with siRNAs that target METTL3 and METTL14 as well as negative controls without siRNA (mock), with scrambled siRNA and siRNAs that target Cyclophilin B. RNA expression of METTL3 (blue) and METTL14 (red) was normalized to mock transfection. Error bars represent standard deviation. Data shown are biological replicates of Fig 1C. **(C)** Two rounds of polyadenylated RNA selection (top) were performed for each siRNA condition in (C). Decreasing amounts of RNA from each condition was probed to detect the m^6^A modification. Total RNA was isolated from each siRNA condition (bottom). The RNA was depleted of rRNA and treated with RNase R to digest linear RNAs. Decreasing amounts of RNA from each condition was probed to detect the m^6^A modification (bottom). Controls were performed as described in Fig S1C.

**Figure S2, Related to Figure 2:**
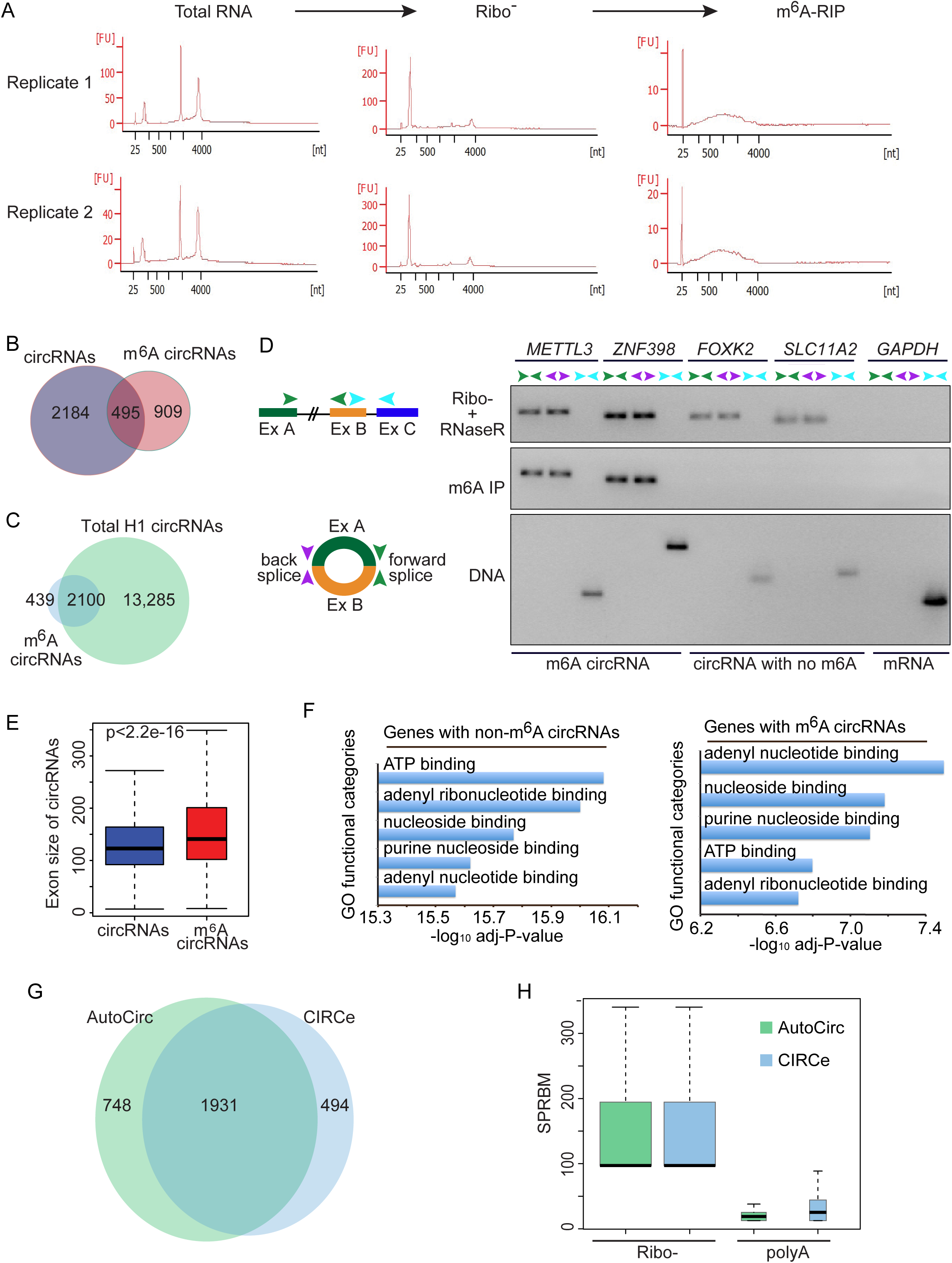
m^6^A-circRNAs identified in human embryonic stem cells. **(A)** Bioanalyzer analysis of RNA in each step of sample preparation prior to library construction. **(B)** Venn diagram showing the total circRNAs identified from the input sample (Ribo-RNA) and m^6^A-circRNAs identified after m^6^A IP. **(C)** Venn diagram showing the total circRNAs identified from all hESC samples (including our input and ENCODE non-polyA RNA) and m^6^A-circRNAs identified after m^6^A IP. **(D)** Primers were designed to detect forward splice junctions contained in circRNAs (green), back splice junctions contained in circRNAs (purple) and forward splice junctions that were not contained in circRNAs (light blue). RT-PCR was performed after rRNA depletion (Ribo-) and RNase R treatment of total RNA to enrich for circRNAs, after Ribo- and RNase R treatment followed by m^6^A IP to enrich for m^6^A-circRNAs, and on genomic DNA. The primers used to detect splice junctions that were not contained in circRNAs (light blue) were amplified from genomic DNA to confirm that the primers are functional. GAPDH does not encode circRNAs and serves as a negative control. The primers designed to amplify forward splices contained in circRNAs (green) were too distant to be amplified from genomic DNA. **(E)** Comparison of exon size of total circRNAs and m^6^A-circRNAs. **(F)** The most enriched GO molecular functional categories of parent genes of non-m^6^A-circRNAs (left) and m^6^A-circRNAs (right). **(G)** The Venn diagram showing the circRNAs identified by our AutoCirc pipeline and the CIRCexplorer (CIRCe) pipeline. About 80% of circRNAs found by CIRCexplorer are also identified by our AutoCirc pipeline. The pools of circRNAs identified by these two methods very similar (p<1.6e-295 by hypergeometric test). **(H)** The expression levels in SPRBM of back splice junctions of circRNAs identified by our AutoCirc computational pipeline and CIRCexplorer (CIRCe) pipeline (Zhang et al., 2014) from our hESC Ribo-data and the polyA data. SRPBM is the back-spliced reads per billion mapping metric, see Materials and Methods for details.) Both our computational pipeline AutoCirc and CiRCexplorer identified more than 2000 circRNAs from Ribominus RNA-seq data with 109million reads whereas both of them identify several hundreds lowly expressed of circRNAs from polyA-selected RNA-seq data with 557 million reads. It indicates that both circRNAs methods have similar level of accuracy and sensitivity of detecting circRNAs.

**Figure S3, related to Figure 3:**
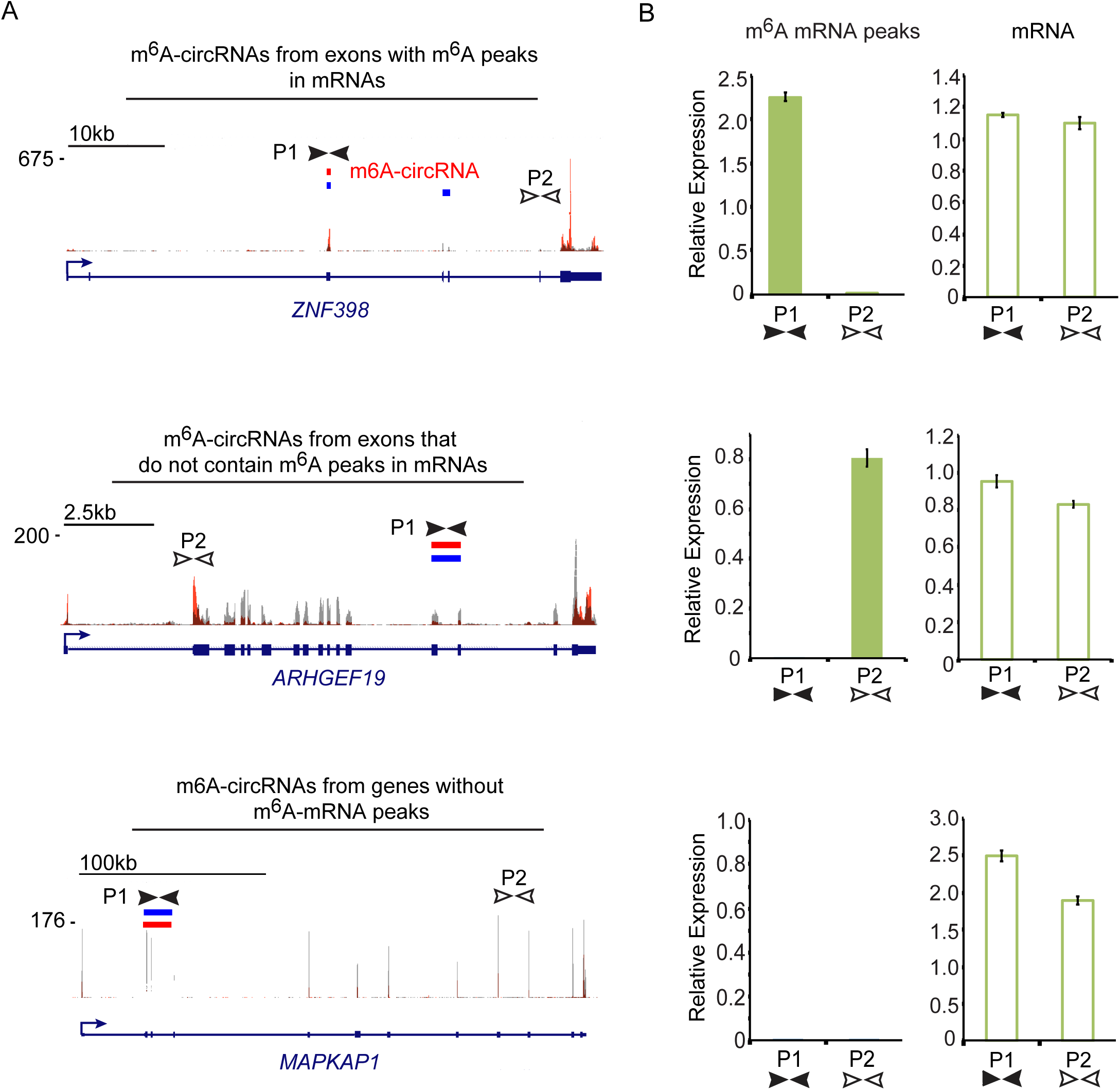
Many m6A-circRNAs contain exons that are not methylated in mRNAs. **(A)** Gene tracks showing the location of m^6^A-circRNAs and m^6^A mRNA peaks in three example genes. *ZNF398* (top) produces an m^6^A-circRNA (red box) from the same exon that contains an m^6^A mRNA peak. *ARHGEF19* (middle) produces an m^6^A-circRNA from an exon that does not contain an m^6^A peak in the mRNA, and the mRNA contains an m^6^A peak in a different exon. *MAPKAP1* (bottom) produces an m^6^A-circRNA, and there is no m^6^A peak in the mRNA produced by the same gene. Red bars indicate m^6^A-circRNAs detected by sequencing. Blue bars represent total circRNAs identified by sequencing. Orange peaks represent m^6^A peaks from identified in mRNAs (Batista et al., 2014) and gray peaks represent mRNA background levels. The structure of each gene is indicated below each track and arrows indicate the direction of transcription. The location of primer sets for each transcript is indicated by black and white arrows (P1 and P2). **(B)** qRT-PCR was performed on cDNA prepared following polyA selection and m^6^A IP (m^6^A mRNA peaks) and following polyA selection alone (mRNA) for the genes in (B). P1 primers amplify m^6^A circRNAs, but only in the top example do the same primers amplify mRNA that contains an m^6^A peak. Amplification of the P2 primers for *ARHGEF19* (middle) indicates that the mRNA contains an m^6^A peak at a different exon. No m^6^A peaks are detected in *MAPKAP1* mRNA (bottom). Each primer is amplified in polyA-selected RNA (right), indicating that all primers can amplify mRNAs before m^6^A IP. These data represent additional examples of what is shown in Figures 3D and 3E.

**Figure S4, related to Figure 5:**
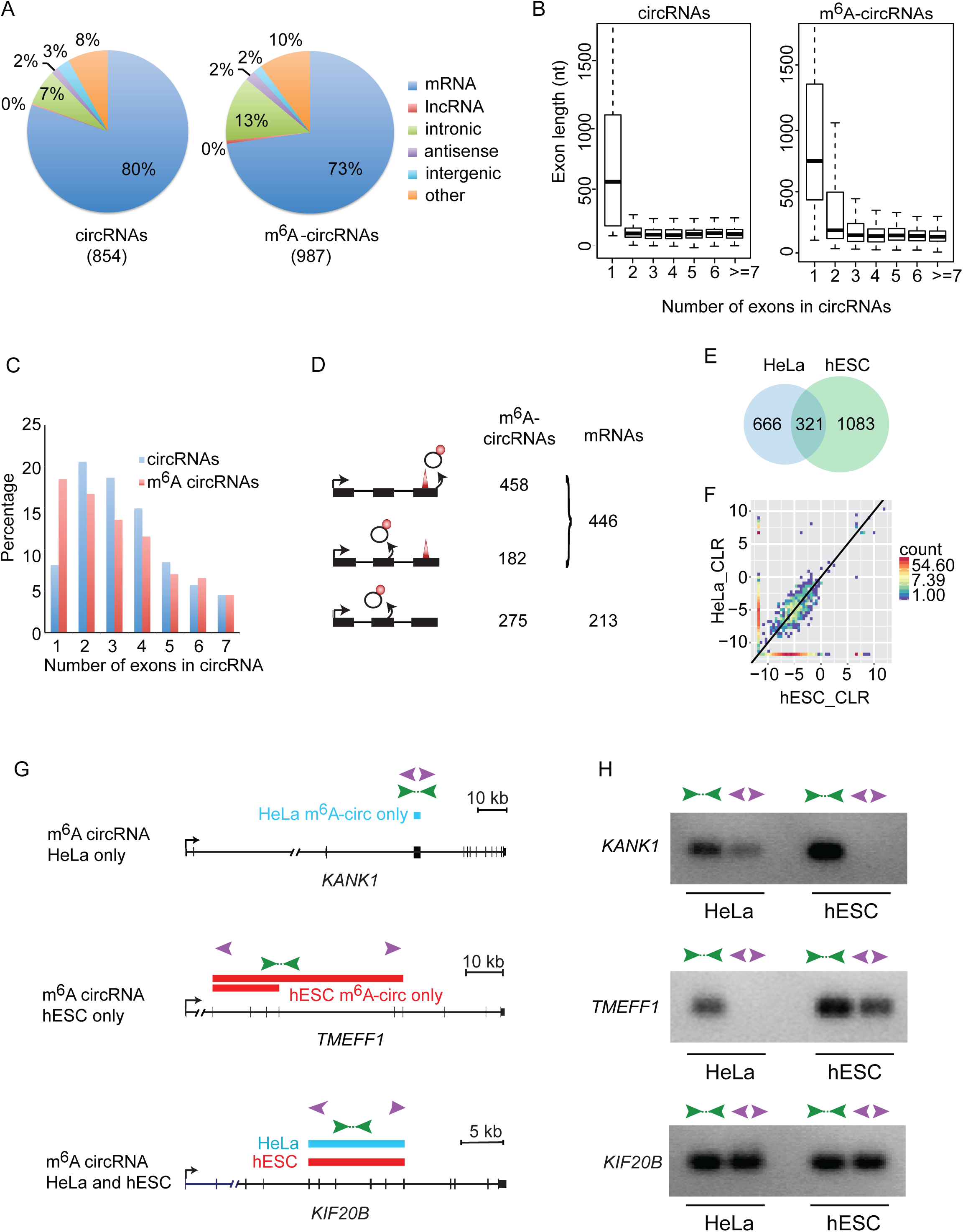
m6A-circRNAs show cell-type-specific patterns of expression. **(A)** The genomic distribution of total circRNAs (left) and m^6^A-circRNAs (right) in HeLa cells. The total number of circRNAs identified in each condition is shown in parenthesis. **(B)** The distribution of exon length (y-axis) for total circRNAs and m^6^A-circRNAs is plotted based on the number of exons spanned by each circRNA (x-axis). **(C)** The number of exons spanned by each total circRNA (blue) and each m^6^A-circRNA (red) was calculated. The percentage of circRNAs (y-axis) spanning up to seven exons (x-axis) is shown. **(D)** The distribution of m^6^A-modified exons between circRNAs and mRNAs in HeLa cells. **(E)** Venn diagram showing the overlap between m^6^A-circRNAs identified in HeLa cells and those identified in hESCs. **(F)** Two-dimensional histogram comparing the expression levels of all m^6^A circRNAs in HeLa cells and these in hESCs. **(G)** Tracks showing additional examples of m^6^A circRNAs that are unique to HeLa cells (top), that are unique to hESCs (middle), and are common between hESC and HeLa cells (bottom). Blue rectangles indicated m^6^A-circRNAs identified in HeLa cells, and red rectangles indicate m^6^A-circRNAs identified in hESCs. Green arrows indicate the location of primers that amplify across forward splice junctions and purple arrows indicate primers that amplify across back splices. **(H)** RT-PCR was performed on RNA prepared following rRNA depletion, RNase R treatment and m^6^A IP in HeLa and hESCs, respectively.

**Figure S5, related to Figure 5:**
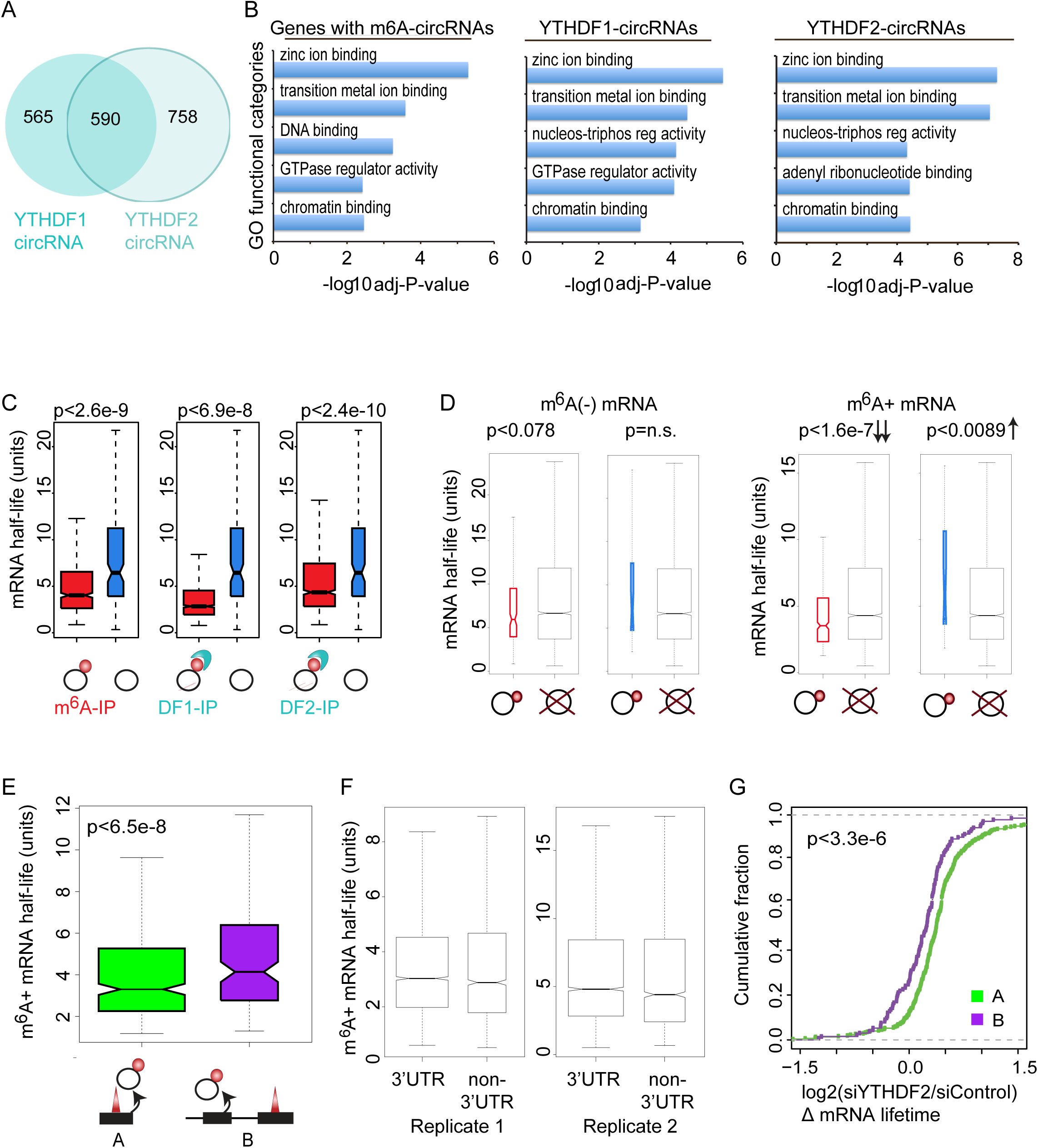
m^6^A-circRNAs bind YTHDF1 and YTHDF2 and identify transcripts with shorter half-lives. **(A)** Venn diagram showing the overlap between circRNAs interacting with YTDHF1 (left) and YTHDF2 (right) in Hela cells. **(B)** The most enriched GO molecular functional categories of the parent genes of m^6^A-circRNAs (left), YTHDF1-bound circRNAs (middle) and YTHDF2-bound circRNAs (right). **(C)** The half-life of mRNAs for parent genes that encode m^6^A circRNAs (left), YTHDF1-bound (DF1-IP) circRNAs (middle) or YTHDF2-bound (DF2-IP) circRNAs (right). Black rings represent circRNAs, red circles indicate m^6^A modification, and light blue structures represent YTH proteins. **(D)** The mRNA half-life for m^6^A negative (-) (left) and m^6^A positive (+) (right) mRNAs whose parent genes encode m^6^A-circRNAs (black ring with red circle attached), circRNAs without m^6^A (black ring) or no circRNAs (black ring with red “X”). The p-values are indicated above each comparison and downward arrows indicate a statistically significant change in half-life. **(E)** The half-life of m^6^A-mRNAs whose parent genes also encode m^6^A-circRNAs from the same exon(s) where an m^6^A peak is found in mRNA (labeled A) is compared to the half-life of m^6^A-mRN whose parent gene encodes an m^6^A circRNA at exon(s) where no m^6^A peak is found in mRNAs (labeled B). The p-value is indicated. **(F)** The half-life was calculated for m^6^A-mRNAs containing m^6^A modifications in the 3’ UTR and m^6^A-mRNAs with modifications outside the 3’UTR for both replicates. **(G)** The change in half-life (log2(siYTHDF2/siControl)) was calculated as described (Wang et al., 2014) for condition A (green) and condition B (purple). The p-values were calculated using Wilcox Mann-Whitney test. Data shown in Fig S5C-S5E, S5G are the results of a second biological replicate of the data shown in Fig 5F–5I.

## References

Batista, P.J., Molinie, B., Wang, J., Qu, K., Zhang, J., Li, L., Bouley, D.M., Lujan, E., Haddad, B., Daneshvar, K., et al. (2014). m6A RNA Modification Controls Cell Fate Transition in Mammalian Embryonic Stem Cells. Cell Stem Cell 15, 707–719.

Caudron-Herger, M., Pankert, T., and Rippe, K. (2016). Regulation of nucleolus assembly by non-coding RNA polymerase II transcripts. Nucleus 7, 308–318.

Chen, C.Y., and Sarnow, P. (1995). Initiation of protein synthesis by the eukaryotic translational apparatus on circular RNAs. Science 268, 415–417.

Chen, K., Lu, Z., Wang, X., Fu, Y., Luo, G.Z., Liu, N., Han, D., Dominissini, D., Dai, Q., Pan, T., et al. (2015). High-resolution N6-methyladenosine (m6A) map using photo-crosslinking-assisted m6A sequencing. Angew. Chemie - Int. Ed. 54, 1587–1590.

Consortium, T.E.P. (2012). An integrated encyclopedia of DNA elements in the human genome. Nature 489, 57–74.

Dominissini, D., Moshitch-Moshkovitz, S., Schwartz, S., Salmon-Divon, M., Ungar, L., Osenberg, S., Cesarkas, K., Jacob-Hirsch, J., Amariglio, N., Kupiec, M., et al. (2012). Topology of the human and mouse m6A RNA methylomes revealed by m6A-seq. Nature 485, 201–206.

Guo, L., and Shorter, J. (2015). It’s Raining Liquids: RNA Tunes Viscoelasticity and Dynamics of Membraneless Organelles. Mol. Cell 60, 189–192.

Hansen, T.B., Jensen, T.I., Clausen, B.H., Bramsen, J.B., Finsen, B., Damgaard, C.K., and Kjems, J. (2013). Natural RNA circles function as efficient microRNA sponges. Nature 495, 384–388

Hansen, T.B., Venø, M.T., Damgaard, C.K., and Kjems, J. (2015). Comparison of circular RNA prediction tools. Nucleic Acids Res. 44, 1–8.

Holley, R.W., Apgar, J., Everett, G.A., Madison, J.T., Marquisee, M., Merrill, S.H., Penswick, J.R., and Zamir, A. (1965). Structure of a Ribonucleic Acid. Science (80-.). 147.

Jeck, W.R., Sorrentino, J.A., Wang, K.A.I., Slevin, M.K., Burd, C.E., Liu, J., Marzluff, W.F., and Sharpless, N.E. (2013). Circular RNAs are abundant, conserved, and associated with ALU repeats. RNA 19, 141–157.

Jiang, L., Zhang, J., Wang, J.-J., Wang, L., Zhang, L., Li, G., Yang, X., Ma, X., Sun, X., Cai, J., et al. (2013). Sperm, but not oocyte, DNA methylome is inherited by zebrafish early embryos. Cell 153, 773–784.

Ke, S., Alemu, E.A., Mertens, C., Gantman, E.C., Fak, J.J., Mele, A., Haripal, B., Zucker-Scharff, I., Moore, M.J., Park, C.Y., et al. (2015). A majority of m6A residues are in the last exons, allowing the potential for 3’ UTR regulation. Genes Dev. 29, 2037–2053.

Kelley, D.R., Hendrickson, D.G., Tenen, D., and Rinn, J.L. (2014). Transposable elements modulate human RNA abundance and splicing via specific RNA-protein interactions. Genome Biol. 15, 537.

Langmead, B., and Salzberg, S.L. (2012). Fast gapped-read alignment with Bowtie 2. Nat. Methods 9, 357–359.

Lasda, E., and Parker, R. (2016). Circular RNAs Co-Precipitate with Extracellular Vesicles: A Possible Mechanism for circRNA Clearance. PLoS One 11, 1–11.

Lin, Y., Protter, D.S.W., Rosen, M.K., and Parker, R. (2015). Formation and Maturation of Phase-Separated Liquid Droplets by RNA-Binding Proteins. Mol. Cell 60, 208–219.

Linder, B., Grozhik, A. V, Olarerin-George, A.O., Meydan, C., Mason, C.E., and Jaffrey, S.R. (2015). Single-nucleotide-resolution mapping of m6A and m6Am throughout the transcriptome. Nat. Methods 12, 767–772.

Liu, J., Yue, Y., Han, D., Wang, X., Fu, Y., Zhang, L., Jia, G., Yu, M., Lu, Z., Deng, X., et al. (2014). A METTL3-METTL14 complex mediates mammalian nuclear RNA N6-adenosine methylation. Nat. Chem. Biol. 10, 93–95.

Marzahn, M.R., Marada, S., Lee, J., Nourse, A., Kenrick, S., Zhao, H., Ben-Nissan, G., Kolaitis, R.-M., Peters, J.L., Pounds, S., et al. (2016). Higher-order oligomerization promotes localization of SPOP to liquid nuclear speckles. EMBO J. 35, 1–22.

Memczak, S., Jens, M., Elefsinioti, A., Torti, F., Krueger, J., Rybak, A., Maier, L., Mackowiak, S.D., Gregersen, L.H., Munschauer, M., et al. (2013). Circular RNAs are a large class of animal RNAs with regulatory potency. Nature 495, 333–338.

Meyer, K.D., Saletore, Y., Zumbo, P., Elemento, O., Mason, C.E., and Jaffrey, S.R. (2012). Comprehensive Analysis of mRNA Methylation Reveals Enrichment in 3’ UTRs and near Stop Codons. Cell 149, 1635–1646.

Meyer, K.D., Patil, D.P., Zhou, J., Zinoviev, A., Skabkin, M.A., Elemento, O., Pestova, T.V., Qian, S.-B., and Jaffrey, S.R. (2015). 5’ UTR m(6)A Promotes Cap-Independent Translation. Cell 163, 999–1010.

Mishima, E., Jinno, D., Akiyama, Y., Itoh, K., Nankumo, S., Shima, H., Kikuchi, K., Takeuchi, Y., Elkordy, A., Suzuki, T., et al. (2015). Immuno-northern blotting: Detection of RNA modifications by using antibodies against modified nucleosides. PLoS One 10, 1–17.

Polioudakis, D., Abell, N.S., and Iyer, V.R. (2015). miR-503 represses human cell proliferation and directly targets the oncogene DDHD2 by non-canonical target pairing. BMC Genomics 16, 40.

Quinlan, A.R., and Hall, I.M. (2010). BEDTools: a flexible suite of utilities for comparing genomic features. Bioinformatics 26, 841–842.

Rybak-Wolf, A., Stottmeister, C., Glažar, P., Jens, M., Pino, N., Giusti, S., Hanan, M., Behm, M., Bartok, O., Ashwal-Fluss, R., et al. (2015). Circular RNAs in the Mammalian Brain Are Highly Abundant, Conserved, and Dynamically Expressed. Mol. Cell 1–16.

Salzman, J., Gawad, C., Wang, P.L., Lacayo, N., and Brown, P.O. (2012). Circular RNAs are the predominant transcript isoform from hundreds of human genes in diverse cell types. PLoS One 7, e30733.

Suzuki, H., Zuo, Y., Wang, J., Zhang, M.Q., Malhotra, A., and Mayeda, A. (2006). Characterization of RNase R-digested cellular RNA source that consists of lariat and circular RNAs from pre-mRNA splicing. Nucleic Acids Res. 34, e63.

Wang, Y., and Wang, Z. (2015). Efficient backsplicing produces translatable circular mRNAs. RNA 21, 172–179.

Wang, P., Doxtader, K.A., and Nam, Y. (2016a). Structural Basis for Cooperative Function of Mettl3 and Mettl14 Methyltransferases. Mol. Cell 63, 306–317.

Wang, X., Lu, Z., Gomez, A., Hon, G.C., Yue, Y., Han, D., Fu, Y., Parisien, M., Dai, Q., Jia, G.,et al. (2014a). N6-methyladenosine-dependent regulation of messenger RNA stability. Nature 505, 117–120.

Wang, X., Zhao, B.S., Roundtree, I.A., Lu, Z., Han, D., Ma, H., Weng, X., Chen, K., Shi, H., and He, C. (2015). N6-methyladenosine Modulates Messenger RNA Translation Efficiency. Cell 161, 1388–1399.

Wang, X., Feng, J., Xue, Y., Guan, Z., Zhang, D., Liu, Z., Gong, Z., Wang, Q., Huang, J., Tang, C.,et al. (2016b). Structural basis of N6-adenosine methylation by the METTL3–METTL14 complex. Nature 534, 1–15.

Wang, Y., Li, Y., Toth, J.I., Petroski, M.D., Zhang, Z., and Zhao, J.C. (2014b). N(6)- methyladenosine modification destabilizes developmental regulators in embryonic stem cells. Nat. Cell Biol. 16, 1–10.

Zhang, H., Elbaum-Garfinkle, S., Langdon, E.M., Taylor, N., Occhipinti, P., Bridges, A.A., Brangwynne, C.P., and Gladfelter, A.S. (2015). RNA Controls PolyQ Protein Phase Transitions. Mol. Cell 60, 220–230.

Zhang, X.-O., Wang, H.-B., Zhang, Y., Lu, X., Chen, L.-L., and Yang, L. (2014). Complementary Sequence-Mediated Exon Circularization. Cell.

Zhou, J., Wan, J., Gao, X., Zhang, X., Jaffrey, S.R., and Qian, S. (2015). Dynamic m 6 A mRNA methylation directs translational control of heat shock response.

